# The impact of Fisher’s Reproductive Compensation on raising equilibrium frequencies of semi-dominant, non-lethal mutations under mutation/selection balance

**DOI:** 10.1101/2021.11.19.469230

**Authors:** Ian M. Hastings

## Abstract

Fisher’s reproductive compensation (fRC) occurs when a species’ demography means the death of an individual allows increased survival of his/her relatives, usually assumed to be full sibs. This likely occurs in many species, including humans. Several important recessive human genetic diseases cause early foetal/infant death allowing fRC to act on these mutations. The impact of fRC on these genetic conditions has been calculated and shown to be substantial as quantified by ω, the fold increase in equilibrium frequencies of the mutation under fRC compared to its absence i.e. ω=1.22 and ω=1.33 for autosomal and sex-linked loci, respectively. However, the impact of fRC on the frequency of the much large class of semi-dominant, non-lethal mutations is unknown. This is calculated here by a mixture of simulation and algebra and shown that ω=2-*h*\**s* and ω≈2-0.19s-0.85h*s for autosomal and sex-linked loci respectively where *h* is dominance (varied between 0.05 and 0.95) and *s* is selection coefficient (varied between 0.05 and 0.9). These results show that the actions of fRC can almost double equilibrium frequency of mutations with low values of *h* and/or *s*. The dynamics of fRC acting on this type of mutation are also identified and discussed.

## 1. Introduction

Population genetics as a discipline started in the 1930s and, at that time, was largely concerned with how advantageous mutations could spread through populations (positive selection) and how deleterious mutations could be eliminated (negative selection). The original assumptions underlying the calculations were simple: random mating of parental genotypes, non-overlapping generations, and independent fates of parents and offspring ([1, 2]). This simple approach has been astonishingly successful and influential over the last 100 years and has been adapted to readily incorporate relaxations to these assumptions such as non-random mating through population sub-structuring, the effects of non-overlapping generations, genetic substructure and “inclusive fitness”. One relaxation, started by R.A. Fisher in 1949, was the realisation that in some species (notably humans) the fates of offspring were not independent and the death of one individual was compensated, to various degrees, by increased survival of its siblings. This occurs in humans because, prior to the demographic transition, females reproduced approximately every 4 to 5 years and early death of a foetus/infant allows the mother to re-conceive earlier and produce a “replacement” sibling and, post demographic transition, human females may tend to choose a family size and “compensate” for any deaths by producing “replacement” siblings. The demographies of many other, non-human species also contain periods of intense intra-brood competition in which death of an individual likely results in increased survival of his/her siblings (see [3] for a recent application to plants which also considered the impact of fRC on the evolution of mating systems). I term this effect Fisher’s reproductive compensation, fRC, to distinguish it from a more recent form of ‘adaptive’ reproductive compensation (i.e. facultatively increasing reproductive effort in animals mating with a sub-optimal mate (e.g. [4]). The impact of fRC has been studied in relation to the obvious example of lethal human genetic diseases. As would be expected intuitively such compensation reduces selection against the recessive lethal allele (assuming the mutation is recessive lethal, the “replacement” siblings have a 2/3 chance of carrying the allele) and the equilibrium frequency of the lethal mutation therefore increases substantially (by around 22% to 33% for autosomal and sex-linked loci respectively e.g. [5]). The impact of fRC therefore has implications for the “genetic load” carried by a population. In the example of recessive lethals, it means an increased proportion of affected individuals in the population (at least at fertilisation). This has social implications with some commentators, including the well know population geneticist Jim Crow, asking how relaxed selection in modern human societies will translate into increased survival of genetically compromised individuals and hence into an increased genetic load [6].

Genetic calculations for the impact of reproductive compensation on recessive lethals alleles, both autosomal and sex-linked, have been developed over the last 70 years, but to date, no such calculations have been made for semi-dominant, non-lethal alleles. and this remains a significant knowledge gap. This manuscript describes how these calculations can be made and quantifies the possible impact of fRC in increasing the equilibrium frequency of such alleles.

## 2. Methods

### 2.1 Biological considerations and general modelling strategy

I adopt the nomenclature of “+” denoting the wildtypes allele and’*m*’ denoting the deleterious mutant allele, giving three autosomal genotypes ++, +*m*, and *mm*. In the case of sex-linked loci, males carry only a single allele plus the Y chromosome giving 3 female genotypes ++, +*m*, and *mm* and two male diploid genotypes +Y and *m*Y. As in conventional population genetics, the frequency of the wildtype allele is denoted *p* and frequency of the mutant is *q*. Fitness of the ++ genotype is 1, fitness of +*m* is 1-*h*s* and of *mm* is 1-*s* where *h* is dominance and *s* is the selection coefficient acting against the double mutant genotype. When considering sex-linked loci the fitness of the male +Y genotype is 1 and that of the *m*Y is 1-*s*. Mutation is assumed to occur at the gamete stage and occurs from wildtype to mutant (“back mutation” from mutant to wildtype is assumed to be negligible).

Previous models of fRC considered recessive lethals so only two adult autosomal genotypes were present i.e. ++ and +*m*. fRC could therefore only operate in mating involving +*m* by +*m* genotypes where 25% of offspring would die and the “replacement” genotypes ++ and +*m* each had fitness of 1 (because *h*=0). The standard way to allow for fRC in this case was to calculate brood size, *B*, after fRC had occurred, as follows

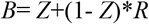

Where *Z* is the proportion of the brood surviving genetic selection, and (1-*Z*) is the proportion dying. Setting *R*=0 indicates that fRC is absent, while setting R=1 restores full brood size.

The problem when considering semi-dominant mutations is that different values of R may occur within different broods. For example, if *s*=0.95 (so 95% of *mm* genotypes die) then in a *mm* by *mm* matin, 95% of the brood may die, and the ability of fRC to restore the full brood size may be implausible in this brood, but capable of fully restoring brood size in a ++ by +*m* mating where a maximum of 50% of offspring may die. This could be addressed by using different R value for each mating types e.g. R may be close to 1 in mating of ++ with +*m* but may be much lower in mating of *mm* with *mm* where most offspring may die. Here, an alternative, more explicit approach is developed based on the potential for “competitive release”, CR, within the brood (the phrase is borrowed from malaria intra-host dynamics [7] which recognises that death of some individuals frees up resources that allows increased survival of the remaining malaria parasites competing for those resources; the same phrase is used in ecology to describe competition between species rather than individuals). Competitive release within broods can be expressed as

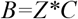

Where C is the competitive release coefficient. The baseline value of C=1 means competitive release is absent, a value of C=2 states that survival of individuals in a reduced brood size can potentially double due to competitive release, and so on. Importantly, the size of brood after the actions of fRC cannot exceed the value of the brood produced by ++ by ++ mating whose value is set to 1, so B will be the lower of

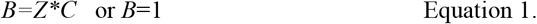

An illustrative example for C=1.5 is given on Figure 1. More sophisticated quantifications of fRC can presumably be constructed to make size of brood after fRC a more complex function of brood size after genetic selection, but the methodology described below can easily incorporate such functions.

**Figure 1.**
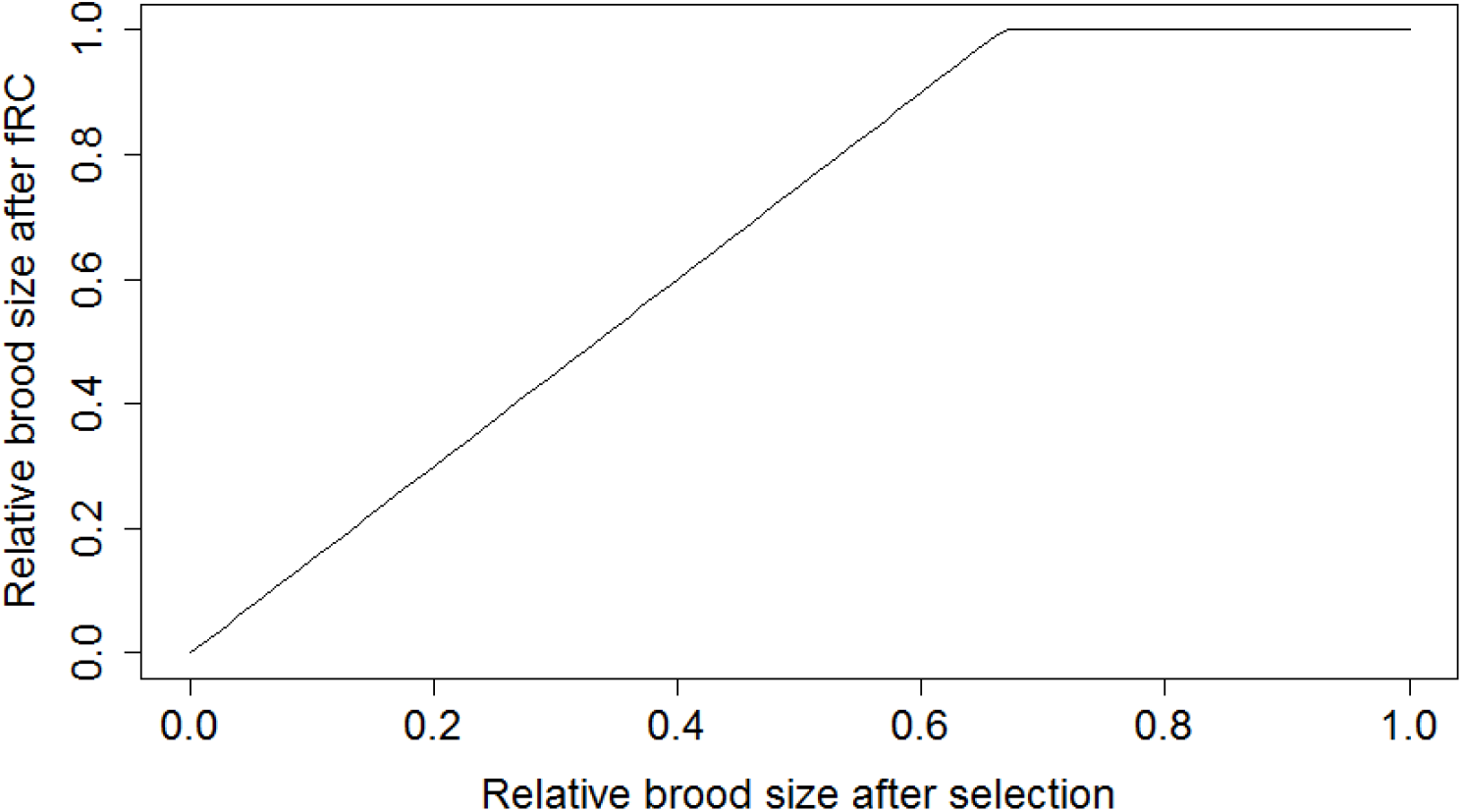
The effects of fRC on brood size after genetic selection. The competitive release coefficient is assumed to be 1.5 and the plot is obtained from Equation 1. Brood sizes after fRC are relative to a brood consisting solely of wildtype homozygotes (whose value is set to 1). In this example fRC can compensate for the loss of up to a third of the brood before its size starts to diminish i.e. when the Xaxis value falls below 0.67.

We assume that fRC occurs within broods of full sibs. It is not necessary to specify the mechanism by which fRC occurs, simply that it exists (for example it may be that, in humans, mother produce an extra offspring to compensate for a dead embryo/foetus/offspring, while in other species it may be intense competition within the brood for resources such as food that leads to fRC).

In standard theory the frequencies of genotypes are tracked independently. However, under fRC the fate of genotypes differ depending on what type of brood they are boorn into e.g. a +*m* genotype may occur in a brood consisting of all +*m* siblings (i.e. from a ++ by *mm* mating), or with a mixture of 50% +*m* and 50% ++ siblings (i.e. from a +*m* by ++ mating). It is therefore necessary to track mating between parental genotypes to recognise that broods differ in genetic composition (as in in Koeslag & Schach [8] although they only required 3 matings as *mm* genotypes are lethal).

The genetic calculations require a multi-stage computation for each of the six mating types, *i*, and three parameters are calculated for each mating type,:

- The frequency of the mating type, denoted *M*_*i*_
- The size of the brood after genetic selection and fRC (from Equation 1), denoted *B*_*i*_
- The proportion of each type of gamete transmitted i.e. *P*_*i*_ for the proportion of wildtype gametes transmitted from the brood and *Q*_*i*_ (=1-*P*_*i*_) for the proportion of mutant alleles

Multiplying these three factors together generates the gametic outputs from each mating type which can then be summed and normalised to obtain gamete frequencies to generate the parental genotypes of the next generation i.e.

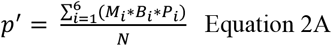

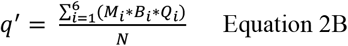

where *N* is the normalising factor i.e.

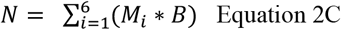

This describes the process for autosomal loci. An analogous process is used when investigating sex-linked loci except that it is necessary to track male and female gametes separately.

The next step is to allow mutation to occur so overall proportion of each type of allele in the population after mutation is

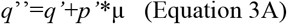

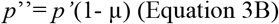

where µ is the mutation rate from wildtype to mutant allele.

These frequencies *p*’’ and *q*’’ are used to calculate the frequency of mating types next generation assuming random union of gametes, and the process iterated until equilibrium is reached

The algorithms were constructed for autosomal and sex-linked loci as described below, then encoded as a R function and run to equilibrium.

### 2.2 Autosomal loci

There are 3 genotypes and hence six different mating combinations. The values of *M*_*i*,_ *B*_*i*,_ *P*_*i*_ and *Q*_*i*_ can be obtained from each mating combination as given below.

*Mating type 1: ++ by ++*

Frequency of this mating types is M_1_= *p*^2*^*p*^2^ _=_ *p*^4^

[Brood genotypes: 100% ++]

Size of brood after selection is Z_1_ = 1

Size of brood after fRC is B_1_=1

Proportion of wildtype allele transmitted from the brood is

P_1_= 1

Proportion of mutant alleles is

Q_1_ = 0

*Mating type 2: ++ with +m*.

Frequency of this mating types is M_2_= 2\**p*^2*^2*pq* =4*p*^3^*q*

[Brood genotypes: 50% ++, 50% +*m*]

Size of brood after selection is Z_2_ = 0.5+0.5(1-*h*s*)

Size of brood after fRC is B_2_=Z_2_*C or B_2_=1 whichever is the lower

Frequencies of genotypes in broods after selection is (f)

*f*(++) = 0.5/*N*

*f*(+*m*) = 0.5 * (1 ™ *h* * *s*)/*N*

where *N* is a normalising factor equal to the sum of the numerators.

Proportion of wildtype alleles transmitted from the brood is

*P*_2_ = [*f*(+ +) + *f*(+*M*) * 0.5]/*W*

And proportion of mutant alleles is

*Q*_2_ = [*f*(+*m*) * 0.5]/*W*

Where W is a normalising factor i.e. W =P_2_+Q_2_

*Mating type 3: ++ with mm*.

Frequency of this mating type is M_3_= 2\**p*^2*^*q*^2^

[Brood genotypes: 100% +m]

Size of brood after selection is Z_3_ = (1-*h*s*)

Size of brood after fRC is B_3_=Z_3_*C or B_3_=1 whichever is the lower

Proportion of wildtype alleles transmitted from the brood is P_3_= 0.5

And proportion of mutant alleles is Q_3_ = 0.5

*Mating type 4: +m with +m*.

Frequency of this mating types is M_4_= 2*pq*\*2*pq* = 4*p*^2^*q*^2^

[Brood genotypes: 25% ++, 50% +*m*, 25% *mm*]

Size of brood after selection is Z_4_ = 0.25+0.5(1-*hs*)+0.25(1-*s*)

Size of brood after fRC is B_4_=Z_4_*C or B_4_=1 whichever is the lower

Frequencies of genotypes in broods after selection is (f)

*f*(++) = 0.25/*N*

*f*(+*m*) = 0.5 * (1 ™ *hs*)/*N*

*f*(*mm*) = 0.25 * (1 ™ *s*)/*N*

where N is a normalising factor equal to the sum of the numerators.

Proportion of wildtype alleles transmitted from the brood is

*P*_4_ = [*f*(+ +) + *f*(+*m*) * 0.5]/*W*

And proportion of mutant alleles is

*Q*_4_ = [*f*(+*m*) * 0.5 + *f*(*mm*)]/*W*

Where W is a normalising factor i.e. W =P_4_+Q_4_

*Mating type 5: +m with mm*.

Frequency of this mating types is M_5_= 2*2*pq*\**q*^2^ = 4*pq*^3^ [Brood genotypes: 50% +*m*, 50% *mm*]

Size of brood after selection is Z_4_ = 0.5(1-*hs*)+0.5(1-*s*)

Size of brood after fRC is B_5_=Z_5_*C or B_5_=1 whichever is the lower

Frequencies of genotypes in broods after selection is (f)

*f*(+*m*) = 0.5 * (1 ™ *hs*)/*N*

*f*(*mm*) = 0.5 * (1 ™ *s*)/*N*

where N is a normalising factor equal to the sum of the numerators.

Proportion of wildtype alleles transmitted from the brood is

*P*_5_ = [*f*(+*m*) * 0.5]/*W*

And proportion of mutant alleles is

*Q*_5_ = [*f*(+*m*) * 0.5 + *f*(*mm*)]/*W*

Where W is a normalising factor i.e. W =P_5_+Q_5_

*Mating type 6: mm with mm*.

Frequency of this mating types is M_6_= *q*^2*^*q*^2^ = *q*^4^

[Brood genotypes: 100% mm]

Size of brood after selection is Z_6_ = (1-*s*)

Size of brood after fRC is B_6_=Z_6_*C or B_6_=1 whichever is the lower

Proportion of wildtype alleles transmitted from the brood is

P_6_= 0

And proportion of mutant alleles is

Q_6_ = 1

These values of M_i_, B_i_, P_i_ and Q_i_ were then used in Equation 2 as described above. Equilibrium frequencies at autosomal loci were obtained using the R function “uniiroot” as only a single variable has to be solved (i.e. equilibrium frequency of mutations).

### 2.3. Sex-linked genes

We assume a XY system i.e. male are the heterogametic sex. There are five diploid genotypes i.e.

♂(+Y), ♂(+Y)

♀(++), ♀(+*m*) ♀(*mm*)

giving six mating types in total.

There are four types of haploid gametes i.e.

♂(+), ♂(*m*),

♀(+), ♀(*m*),

And we denote their frequencies using ‘f’ as a prefix i.e. f♂(+) is frequency of the ♂(+) gamete

There are six different mating combinations and, as for autosomal loci, the values of *M*_*i*,_ *B*_*i*,_ *P*_*i*_ and *Q*_*i*_ can be obtained from each mating combination as given below.

*Mating type 1: ♂(+Y) with ♀(++)*

Frequency of this mating types is M_1_= f♀(+)* f♂(+) *f♀(+)

[Brood genotypes: 50% ♂(+Y), 50%♀(++)]

Size of brood after selection is Z_1_ = 1

Size of brood after fRC is B_1_=1

Frequencies of genotypes in broods after selection is (f)

f♂(+*Y*)=0.5

f♂(*mY*)=0

f♀(++)=0.5

f♀(+*m*)=0

f♀(*mm*)=0

And proportion of each gamete type transmitted from this mating type is

P_1_♂(+)= f♂(+, Y)

P_1_♂(m)=0

P_1_♀(+)= f♀(++)

P_1_♀(m)=0

*Mating type 2: ♂(+Y) with ♀(+m)*

Frequency of this mating types is M_2_=♀(+)* [f♂(+)* f♀(*m*) + f♂(*m*)* f♀(+)]

[Brood genotypes: 25% ♂(+Y), 25% ♂(*m*Y), 25%♀(++), 25%♀(+*m*)]

Size of brood after selection is Z_2_ = 0.25+0.25*(1-*s*)+0.25+0.25(1-*h*s*)

Size of brood after fRC is B_2_=Z_2_*C or B_2_=1 whichever is the lower

Frequencies of genotypes in broods after selection is (f)

f♂(+Y)=0.25/*N*

f♂(mY)=0.25*(1-*s*)/*N*

f♀(++)=0.25/*N*

f♀(+m)=0.25*(1-*h*s*)/*N*

f♀(mm)=0

where *N* is a normalising factor equal to the sum of the numerators.

Proportion of each gamete type transmitted from this mating type is

P_2_♂(+)= f♂(+Y)

P_2_♂(m)= f♂(*m*Y)

P_2_♀(+)= f♀(++)+ f♀(+*m*)*0.5

P_2_♀(m)= f♀(+*m*)*0.5

*Mating type 3: ♂(+Y) with ♀(mm)*

Frequency of this mating types is M_3_= f ♀(+)* f♂(*m*)* f♀(*m*)

[Brood genotypes: 50% ♂(*m*Y), 50%♀(+*m*)]

Size of brood after selection is Z_3_ = 0.5*(1-*s*)+0.5*(1-*h*s*)

Size of brood after fRC is B_3_=Z_3_*C or B_3_=1 whichever is the lower

Frequencies of genotypes in broods after selection is (f)

f♂(+Y)=0

f♂(mY)=0.5*(1-*s*)/*N*

f♀(++)=0

f♀(+m)=0.5*(1-*h*s*)/*N*

f♀(mm)=0

where *N* is a normalising factor equal to the sum of the numerators.

Proportion of each gamete type transmitted from this mating type is

P_3_♂(+)= 0

P_3_♂(m)= f♂(*m*Y)

P_3_♀(+)= f♀(+*m*)*0.5

P_3_♀(m)= f♀(+*m*)*0.5

*Mating type 4: ♂(mY) with ♀(++)*

Frequency of this mating types is M_4_= f♀(*m*)* f♂(+)* f♀(+) [Brood genotypes: 50% ♂(+Y), 50%♀(+*m*)]

Size of brood after selection is Z_4_ = 0.5+0.5*(1-*h*\**s*)

Size of brood after fRC is B_4_=Z_4_*C or B_4_=1 whichever is the lower

Frequencies of genotypes in broods after selection is (f)

f♂(+Y)=0.5/*N*

f♂(*m*Y)=0

f♀(++)=0

f♀(+*m*)=0.5*(1-*h*\**s*)/*N*

f♀(*mm*)=0

where *N* is a normalising factor equal to the sum of the numerators.

Proportion of each gamete type transmitted from this mating type is

P_4_♂(+)= f♂(+Y)

P_4_♂(m)= 0

P_4_♀(+)= f♀(+*m*)*0.5

P_4_♀(m)= f♀(+*m*)*0.5

*Mating type 5: ♂(mY) with ♀(+m)*

Frequency of this mating types is M_5_= ♀(*m*)* [f♂(+)* f♀(*m*) + f♂(m)* ♀(+)]

[Brood genotypes: 25% ♂(+Y), 25% ♂(*m*Y), 25%♀(+*m*), 25%♀(*mm*)]

Size of brood after selection is Z_5_ = 0.25+0.25*(1-*s*)+0.25*(1-*h*s*)+0.25*(1-*s*)

Size of brood after fRC is B_5_=Z_5_*C or B_5_=1 whichever is the lower.

Frequencies of genotypes in broods after selection is (f)

f♂(+Y)=0.25/*N*

f♂(mY)=0.25*(1-*s*)/*N*

f♀(++)=0

f♀(+m)=0.25*(1-*h*s*)/*N*

f♀(mm)=0.25*(1-*s*)/*N*

where *N* is a normalising factor equal to the sum of the numerators.

Proportion of each gamete type transmitted from this mating type is

P_5_♂(+)= f♂(+Y)

P_5_♂(m)= f♂(*m*Y)

P_5_♀(+)= f♀(+*m*)*0.5

P_5_♀(m)= f♀(+*m*)*0.5+ f♀(*mm*)

*Mating type 6: ♂(mY) with ♀(mm)*

Frequency of this mating types is M_6_= f♀(*m*)* f♂(*m*)* ♀(*m*)

[Brood genotypes: 50% ♂(*m*Y), 50%♀(*mm*)]

Size of brood after selection is Z_6_ = (1-s)

Size of brood after fRC is B_6_=Z_6_*C or B_6_=1 whichever is the lower

Frequencies of diploid genotypes in broods after selection is (f)

f♂(+Y)=0

f♂(*m*Y)=0.5*(1-*s*)/*N*

f♀(++)=0

f♀(+*m*)=0

f♀(*mm*)=0.5*(1-*s*)/*N*

where *N* is a normalising factor equal to the sum of the numerators.

Proportion of each gamete type transmitted from this mating type is

P_6_♂(+)= 0

P_6_♂(m)= f♂(*m*Y)

P_6_♀(+)= 0

P_6_♀(m)= f♀(*mm*)

The algorithm then proceeds as in Equation 2 but tracking female and male gametes separately i.e.:

For male gametes

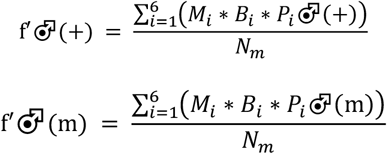

where N_m_ is a normalising factor equal to the sum of the two numerators in the male gamete equations.

For female gametes

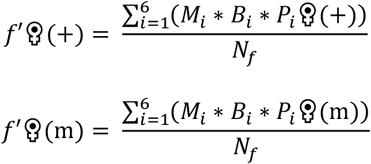

where N_f_ is a normalising factor equal to the sum of the two numerators in the female gamete equations.

Mutation then occurs as before i.e. analogous to Equation 3

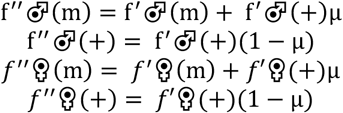

Note that that 2/3 of sex-linked alleles are in females, so overall frequencies of the mutant allele in the population, 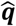, **is**

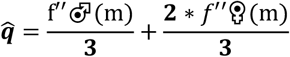

Equilibrium allele frequencies was defined as occurring when mean frequency, 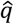 differed by less than a factor of 0.0000001 in consecutive iterations (but see Supplementary material 1 for further discussion). The simulations were started with extremely low frequencies, and extremely high frequencies, and the R scripts verified that both iterate onto the same equilibrium value of 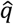 (convergence to equilibrium is rapid for sex-linked loci so this transparent approach was preferred to using the uniroot function).

### 2.4 Sex linked loci with sexual dimorphism

I also investigate the impact of fRC on sex-linked genes in species that are sexually dimorphic to the extent that fRC can only occurs within sexes i.e death of a male can only be compensated by increased survival of his brothers and <u>not</u> by increased survival of his sisters. Similarly for females i.e. death of female can only result in increased survival of her sister. The method used for the simulations were analogous to, and extremely simar with, methods used for sex-linked loci described in 2.3. The only difference is that fRC only occurred with each sex in the brood. The methods are therefore described in Supplementary Material part 2 to avoid repetition.

### 2.5 Checking the algorithm recovers standard equations

Setting CR=1 means fRC is absent and the algorithms should therefore give the standard results for mutation/selection balance given below. Alternatively, if fRC is set high (CR=100 is used here) then the algorithm should obtain the published results on recessive mutations obtained when fRC is sufficiently high that it allows full replacement in all brood types. These previous results are as follows where µ denotes mutation rate:

*Autosomal loci:*

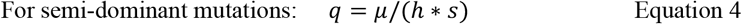

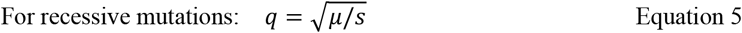

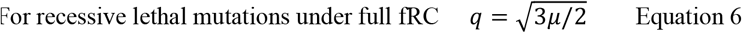

Equations 4 and 5 were as taken from Charlesworth and Charlesworth (their Equations 4.3 and 4.4) and Equation 6 taken Templeton and Yokoyama [5] (noting that and Koeslag and Schach [8] and Hastings [9] give numerically identical versions of Equation 6).

*Sex linked loci:*

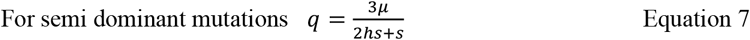

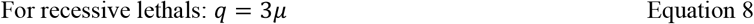

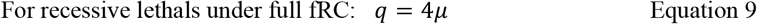

Equation 7 was taken from Charlesworth and Charlesworth (their equation 4.5) and Equations 8 and 9 taken from Templeton and Yokoyama ([5] their equations 4 and 7) and Hastings ([10]; his equations 3 and 4); for simplicity, mutation rates were assumed to be equal in male and females in derivation of all three equations.

### 2.6 Running the simulations

All combinations of the following selection coefficients and dominance values were investigated i.e.

s=0.05, 0.1, 0.15, 0.2….. 0.9

h=0.05, 0.1, 0.15, 0.2….. 0.95

and resulting equilibrium frequency of mutant alleles obtained as described above. Mutation rates were set to be 10^−5^, 10^−6^, 10^−7^, 10^−9^ but did not affect the relative impact of fRC (see later) so a default of 10^−5^ was used.

## 3. Results

### Autosomal loci

The algorithm for autosomal loci reliably recovered the standard, published results in the absence of fRC (Equations 4 and 5) and the published result for recessive lethals when fRC is high (Equation 6).

Figure 2A show the increase in equilibrium frequency under fRC compared to equilibrium frequency its absence, as a function of dominance and selection coefficients. The surface is roughly symmetrical against the dominance and selection which is consistent with expectations as the low equilibrium frequencies of the mutations ensures most mutations will be in the heterozygous form whose fitness is 1-*h*\**s*. The increases were therefore replotted against the value of h*s as shown on Figure 3 for a range of competitive release coefficients. The increase is essentially linear against *h*s* until fRC breaks down and the increase falls relatively rapidly thereafter. The regression coefficients were calculated on the linear portions of the four panels of Figure 3 and had an intercept of 100, with a slope of −100 in all cases, suggesting that the equilibrium frequency in the presence of full fRC can be obtained by updating Equation 2

**Figure 2.**
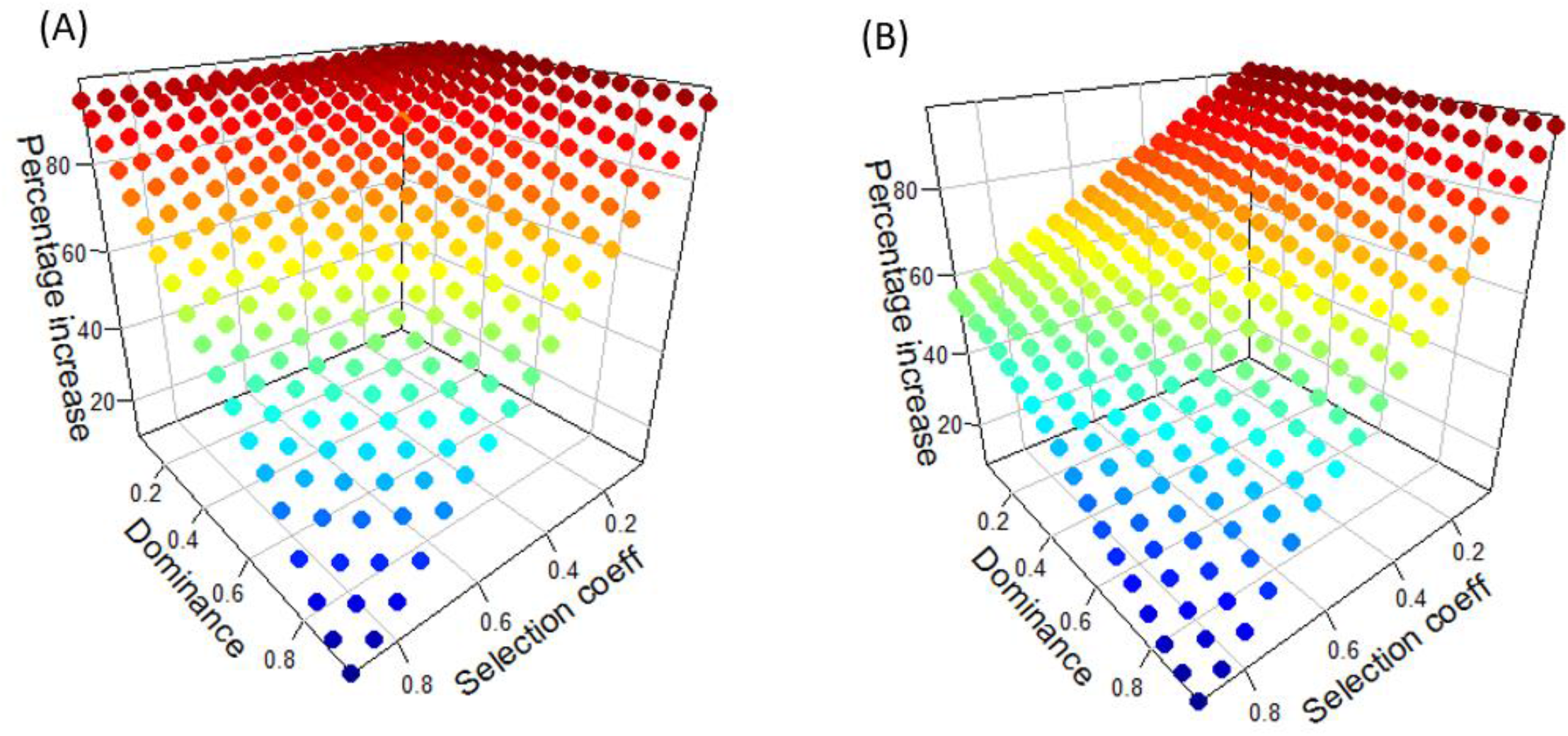
Percentage increase in equilibrium mutant allele frequency attributable to the actions of fRC i.e. compared to standard result in the absence of fRC. Plots show results assuming CR=1.5 and mutation rate is 10^−5^. (A) autosomal loci (B) sex-linked loci.

**Figure 3.**
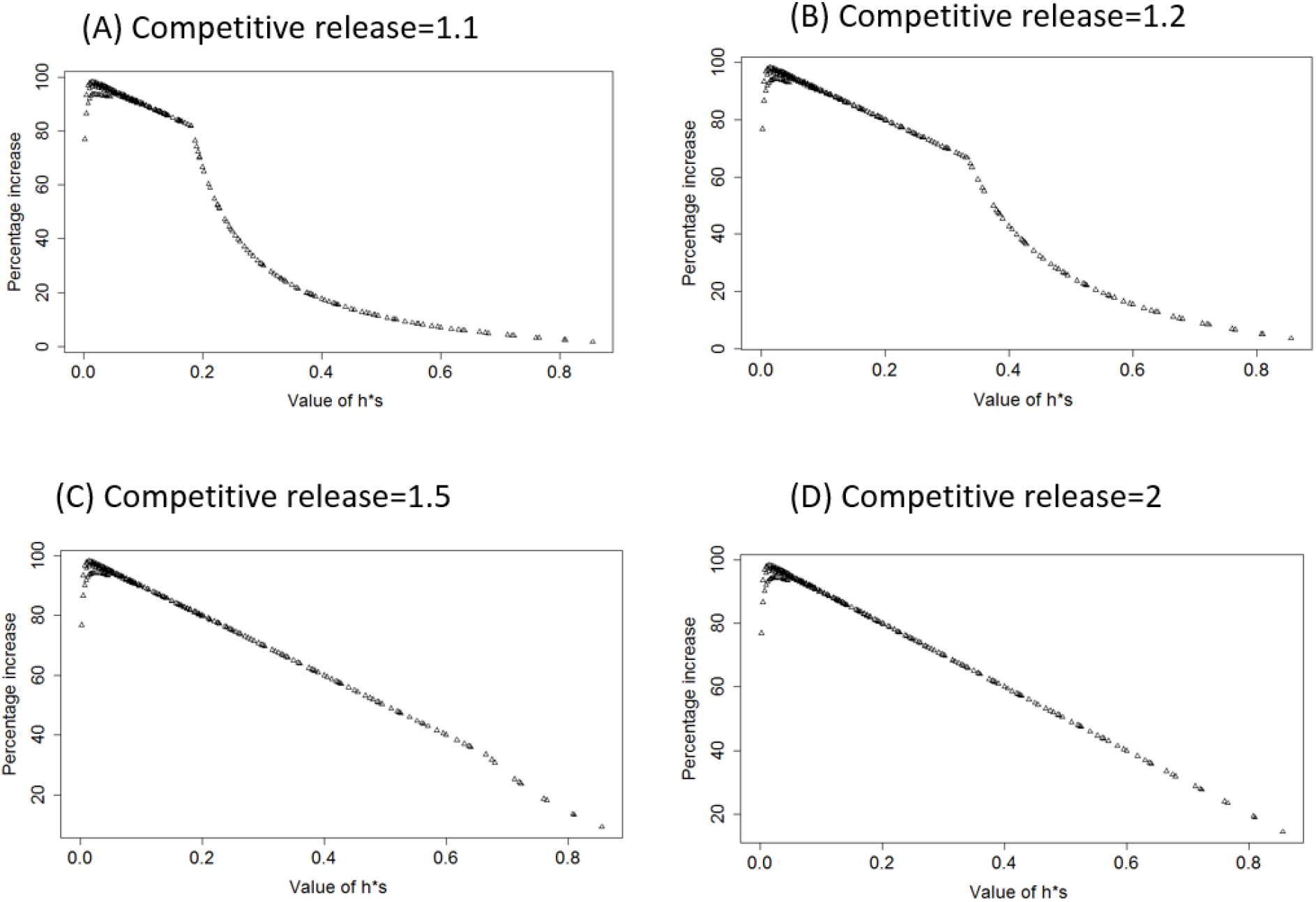
The increase in equilibrium mutant allele frequencies at autosomal loci attributable to fRC i.e. compared to standard theoretical results without fRC. The X axis is *h*s* (i.e. dominance multiplied by selection coefficient) under four illustrative values of competitive release coefficients. Competitive release is unable to fully restore brood size once values of h*s exceed 0.18, 0.34, 0.66 and 1.0 for CR values of 1.1, 1.2, 1.5 and 2, respectively; see Equation 14.

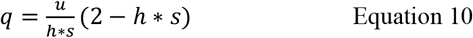

We can recover this result algebraically by noting that when the frequency of mutations is low, most mutant alleles will be in the autosomal mating Type 2: i.e. ++ with +*m* whose brood genotypes are 50% ++ and 50% +m. The fitness of the +m genotypes *w*_*+m*_ in the absence of fRC is therefore

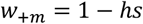

Rising to

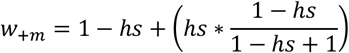

In the presence of fRC. The term in brackets is the number of replacements (“*h*s*” of them) multiplied by the proportion that are of genotype +*m* (the second factor in the brackets). Moving the brackets and simplifying slightly gives

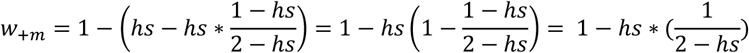

[The last step was obtaining using the algebraic rule that 1-a/b=(b-a)/b which simplifies the expression in brackets to 1/(2-hs)].

This suggest the equilibrium frequency of deleterious mutations under full fRC is

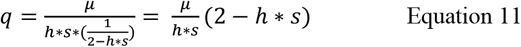

Which is identical to Equation 10 obtained empirically by regression. It holds numerically over the approximate range *h*s*=0.05 to 0.95 provided full fRC occurs (Figure 3, Panel D). This relationship breaks down as h*s becomes very small as selection is then virtually absent, equilibrium frequency rises so the assumption that all selection occurs in autosomal mating type 2 is violated.

The equilibrium frequency in the absence of fRC is u/(h*s) (Equation 4) revealing that that the fold increase in equilibrium frequencies driven by fRC, is

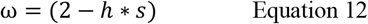

Equations 10 and 11 hold providing the level of CR is sufficient to replace the proportion of individual lost to genetic selection. The critical fraction of the brood, C, that can die and be replaced without a reduction in brood size can be calculated as

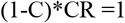

So

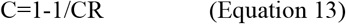

which gives values of C for CR=1.1, 1.2, 1.5 and 2 as 0.09, 0.17, 0.33, and 0.5 respectively. We would therefore expect linearity to break down in Figure 3 when CR is unable to replace all dead brood members. Provided equilibrium mutant allele frequency is sufficiently low that the proportion of double mutant genotypes is negligible, then most offspring with mutant alleles will be in matings ++ by +*m* in which case 50% will be +*m* so mortality in the brood will be 0.5\**h*\**s*. Linearity should therefore break down when

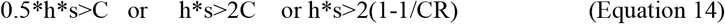

which appears to be the case; see Figure 3.

### Sex-linked loci

The algorithm recovered the standard results in the absence of fRC i.e. Equations 7 and 8. However, the algorithm could not recover the result from Equation 9 i.e. the formula for the equilibrium frequency of a sex-linked recessive lethal under fRC which should be 4u: when CR=2 or 10 the algorithm returns equilibrium frequencies of 4.5u. Extensive bug-checking was performed to try and understand this anomalous result but no computational problems were identified. The value of 4.5u appears to be a valid result for reasons given in the Discussion

Figure 2B provides an example of how fRC increases equilibrium frequency of sex-linked deleterious mutations over a wide parameter range. The increase in mutant allele frequency compared to the standard result (i.e., in the absence of fRC) is far more dependent on selection coefficient than for dominance (compared to autosomal loci). This most likely arises because most selection occurs in males who have only a single copy of the gene meaning dominance has no impact in males. This complicates the plots of the increase against the magnitude of *h*s* shown on Figure 4, compared to the autosomal loci shown on Figure 3, but the same underlying patterns can be discerned i.e. Figure 4 show a steady decline with *h*s* which drops rapidly in Figure 4 panels (a) and (b) as CR become unable to compensate fully for the dead genotypes, with this drop being unnoticeable when CR is 1.5 or 2 (in both Figures 3 and 4) as CR is sufficient to fully compensate for the dead genotypes. The key point is that as h*s becomes small, then the equilibrium frequencies may be double that predicted in the absence of fRC

**Figure 4.**
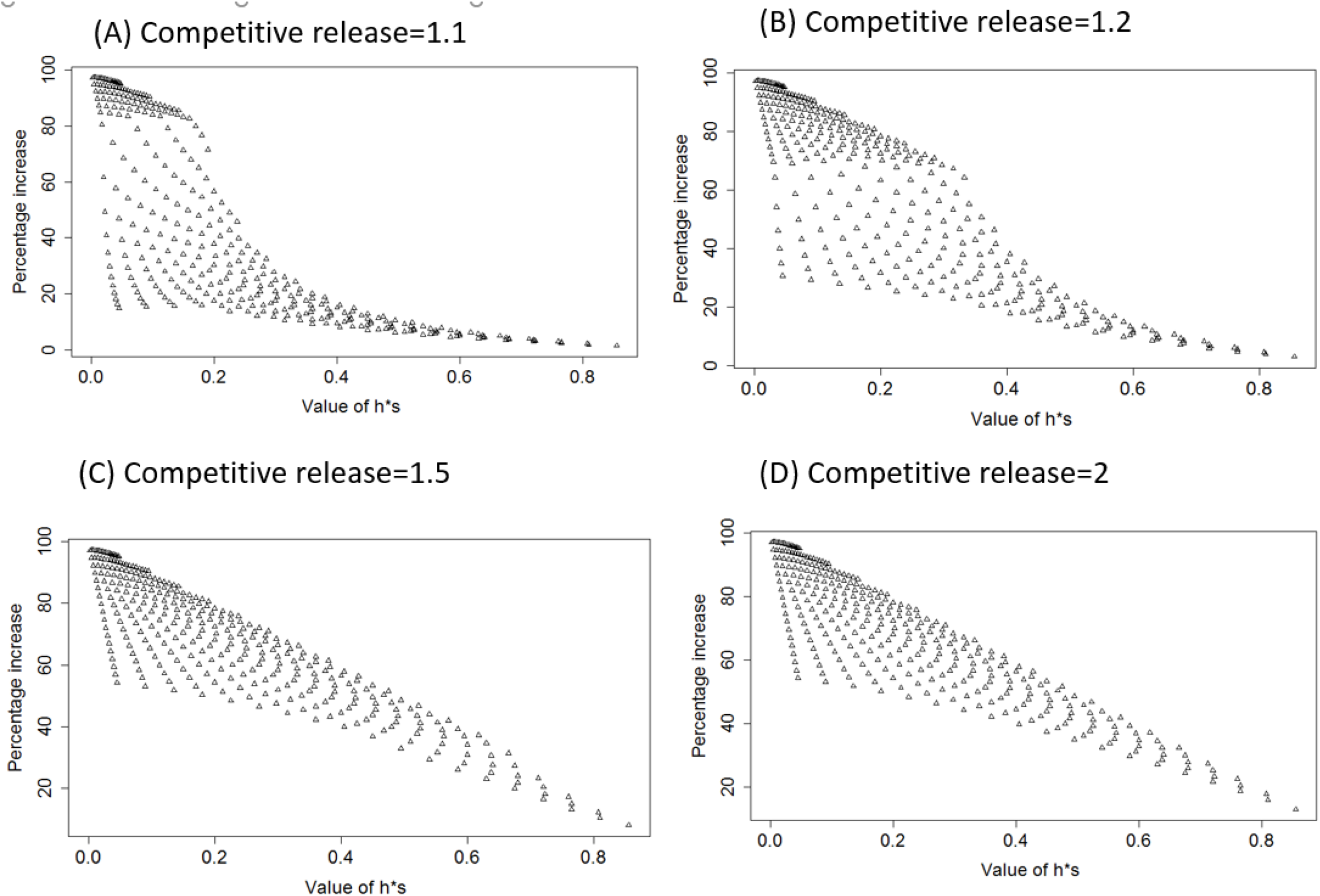
As for Figure 3 but for sex-linked loci.

I could not find an algebraic solution for the increase in equilibrium allele frequency at sex-linked loci caused by fRC but a linear regression of the increases obtained for CR=20 (analogous to the surface shown in Figure 2B) fitting *s, h* and *s*h* returned the empirical result that the fold-increase compared to the standard results driven by fRC, ω is

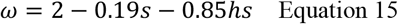

These empirical estimates lie within a proportion 0.97 to 1.04 of the simulation results when CR=20, 2 or 1.5. It starts to break down at lower values of CR when fRC becomes unable to replace the losses at higher values of *h*s*.

### Sex linked loci with sexual dimorphism

The results are presented and discussed in more detail in Supplementary Material #2 but the basic result is that equilibrium allele frequencies can be up to ∼3-fold higher than predicted under standard theory (i.e. in the absence of fRC). Empirically derived estimates of ω obtained by regression were

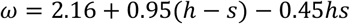

Whose estimates lie within a proportion 0.96 to 1.13 of the simulation results. Slightly more accurate estimates can be made by reducing parameter space to vary dominance from 0.2 to 0.8 and selection coefficient from 0.1 to 0.7 which gives

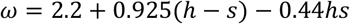

whose values lie within a proportion 0.98 to 1.05 of the simulation values.

## 4. Discussion

The first thing to note is that this work is not directed towards specific mutations (in contrast to previous work that usually investigate recessive lethal mutations of importance to human health), rather it is curiosity-driven to quantify the extent to which life history and demographic traits can result in fRC altering a species genetics and, in particular, the equilibrium frequencies of deleterious mutations and subsequent genetic load. It also establishes a general approach that can recover previous results on mutation/selection balance with, or without, fRC. It lacks algebraic solutions and requires numerical solutions when applied to sex-linked loci, but it is very flexible and can include factors such as inbreeding coefficients into mating frequencies (previous work has shown inbreeding interacts with fRC e.g.[3]) and can potentially include more sophisticated descriptions of fRC compared to the simple equation 1 and Figure 1.

The method recovered all but one of the standard algebraic results already in the literature (i.e., Equations 4 to 9), the exception being Equation 9 i.e. the equilibrium frequency of a sex-linked recessive lethal under fRC where the algorithm retuned a value of 4.5u compared to the published algebraic result of 4u. Extensive bug-hunting failed to identify a clear reason why this discrepancy should exist. The result of 4.5u appears valid for three main reasons: (i) exactly the same code generated the standard result in the absence of fRC (Equation 8), the only difference was in the input parameter used i.e. CR=1 for absence of fRC, CR=2 or 10 for presence of fRC. (ii) The same result arose when the algorithm was recoded to track adult genotype frequencies rather than gametic allelic frequencies (Supplementary Material, part 1). (iii) It seems unlikely that a random bug would produce such a precise result i.e. the value is *exactly* 4.5u. It seems more likely that the discrepancy is due to differing underlying assumptions between the algebraic and computational approaches, but I have been unable to identify this discrepancy. Hence the value of 4.5u stands as an anomalous result but, given the algorithm’s ability to reliable extract the other algebraic result in the literature, it should not threaten the overall quantitative conclusions reached above.

One innovation developed above was to define and use a competitive release coefficient to quantify the extent of fRC (previous models of fRC assumed compensation was fully effective i.e., all brood sizes remained unaffected by genetic deaths). This clarifies the dynamics behind fRC (e.g., Figure 3) which can be explained as the combination of two effects. The first effect is the ability of fRC to compensate for the genetic deaths and maintain the brood size; when the limit of that ability is reached, competitive release become unable to restore full brood size (cf Figure 1), and the breakpoints apparent in the panels of Figure 3 occur. The second effect is that relationship with *h*s* is linear before the breakpoint. Recall that because equilibrium frequency of the mutant is low, the vast majority of the mutations occur in heterozygous form within broods of mainly wildtype genotypes (i.e. in ++ by +*m* matings). The magnitude of *h*s* determines the relative fitness between the mutant heterozygote and their wildtype siblings. As the fitness differences increases, the dead heterozygotes are increasingly replaced by their ++ siblings, hence overall transmission of the mutant from the brood falls as *h*s* increases.

I also investigated the impact of fRC on sex-linked loci when there is complete sexual dimorphism in the life-cycle stages where the fRC is acting i.e. where a dead individual can only be replaced by a survivor of his/her own gender. This is mainly for curiosity and theoretical completeness as I know of no organisms with such strong dimorphism (although I presume this reflects my limited knowledge). There is considerable theoretical work on the impact of differential selection on the sexes on factors such as sex determination, sex ratio, cytonuclear dynamics, etc, so these results are included in the hope that they may be, or may become, helpful in such work. The basic result is fRC in such circumstances can raise equilibrium frequencies of deleterious mutations up to ∼3-fold higher than would occur in the absence of fRC.

In these calculations it was assumed that the fitness penalties <u>only</u> occur during the life-cycle stage(s) in which fRC is acting whereas, in reality, fitness may extend throughout the life. As Porcher & Lande [3] noted in their recent study of fRC and plant mating systems “Much embryo mortality is attributable to early acting, highly deleterious mutations (lethals and semi-lethals), whereas mildly deleterious mutations tend to act late in development during growth”. The calculations above do evaluate semi lethal mutations (generally defined as *s*>0.5, *s*≠1) and I leave it to readers to judge the extent which the assumption of selection only occurring in the presence of fRC is valid. One argument is that the assumption certainly applies to some mutations (i.e. those whose gene expression only occurs early in structured life cycles (such as larvae, pupae, tadpoles, caterpillars) during which fRC may occur. The other, slightly circular argument is that the presence of fRC implies intense selection acting on early stages and these may plausibly dominate later selection pressures. One strategy would be to partition fitness costs between the fRC “brood” and post-fRC “parental” stages and apply them to both periods (Figure S1.1). I did not do so (although it would be relatively straightforward) as the primary purpose of the work was to investigate and close a theoretical gap in quantifying how fRC may potentially affect semi-dominant, non-lethals and present the current work as the limiting case that all selection pressure is occurring in the period when fRC is acting.

Its is clear that fRC is a neglected force in evolution (Hastings unpublished review) whose impact may explain many evolutionary features, not simply increase deleterious allele frequency. This paper has demonstrated computational ways to quantify the impact and given some algebraic descriptions of its impact. In particular the parameters of mutation rate, selection (including dominance) and equilibrium frequencies are all linked in the equations describing mutation/selection balance (MSB; see equations 4 to 9 above) which are useful as estimates of two the factors allow the third to be estimated. The actions of fRC means that application of these equations may introduce up to 2-fold error in calculations. It is important to note that it is not simply a dichotomy between species that do, or do not, allow opportunities for fRC to occur. Probably more important is the dichotomy between genes *within* a species with fRC acting i.e. genes whose expression occurs when fRC is acting and hence is affected by fRC, and those genes whose expression only occurs in life cycle stages after fRC has occurred. For instance, there will be differences in humans between genes whose effects occur early in development (which may be subject to fRC) and genes whose effects occur later after fRC has occurs (such as adult haemoglobins and adult-specific tissues such as eyes, ears). Bioinformatic approaches often compare genes within the same organism and realisation that they may be subject to slightly different selective forces may become important as differences in equilibrium frequencies may be attribute to difference in the magnitude of selection while, in principle, selection pressure acting on the genes may be the same, it is fRC that is driving the differences.

## Supporting information

Supplementary Material, part 1

Supplementary Material, part 2

