## Supplementary Material, part 1 for "The impact of Fisher’s Reproductive Compensation on raising equilibrium frequencies of semi-dominant, non-lethal mutations under mutation/selection balance"

### *Supplementary Material, part 1: Alternative methods of tracking sex-linked loci.*

The simulation in the main text tracks gametic allele frequencies, as is standard in much of population genetics (e.g. [1]). However, simulating gamete production in the brood survivors following fRC inherently means tracking two “generations” per simulation cycle; this is not immediately obvious (at least to me) so is illustrated in Figure S1.1(A). There is only one mutation event and one fRC event in each simulation cycle, so the simulations are valid when run to equilibrium frequencies for autosomes and for sex-linked loci provided mean allele frequency is run to equilibrium (see below).

As part of the process of understanding why tracking sexes separately produced errors for sex-linked loci (see below), and to understand why the simulation did not return Equation 9 in the main text (it returned 4.5μ rather than 4 μ) an alternative algorithm was implemented that tracked diploid genotypes rather than gametes; see Figure S1.1 (B) and the algebraic description below. This eliminated the second generation in the simulation cycle and resulted in one generation per cycle. It returned the same results as the original algorithm based on gametic allele frequencies. Interestingly, for sex-linked loci, the original algorithm used in the main text appeared very slightly more accurate when judged against the standard formula (i.e., Equations 7 and 8 in the main text)

The sex-linked algorithm tracks allele frequencies in male and females separately (because selection is more intense in males), and initial analyses defined equilibrium as being reached when both male and female gametic allele frequencies *tracked separately* were at equilibrium, and the overall equilibrium value, $\hat{q}$ was extracted as the mean frequency as described in the main text (i.e., noting that 2/3 sex-linked genes are in females). Surprisingly, at least to me, this approach did not recover the standard results (i.e., Equations 7 and 8) and recovered results up to 16% higher than the approach taken in the main text (Figure S1.2). The reason seems to be the extra generation per cycle which allows two recombination events per cycle and hence likely equilibrated male and female gametic frequencies. However, if the mean allele frequency across sexes was calculated at the end of each cycle, and the algorithm run to equilibrium mean frequencies, it recovers the standard results currently in the literature (i.e., Equations 7 and 8 in the main text). The results presented in the main text for sex-linked loci were obtained by calculating the mutant allele frequency, combined across males and females, each iteration, and running this value to equilibrium.

In summary, I present the results in the main text based on equilibrating mean allele frequency (Figure S1.1(A)) because they recover standard results, and the purpose of this paper is to address the issue of how fRC affects equilibrium frequencies. I relegate the more arcane matter of simulation structure to this SI. Note also that this choice of equilibrium criterion is conservative i.e. it underestimates the impact of fRC compared to tracking sexes separately.

#### Algorithm for sex-linked genes tracking diploid genotypes to equilibrium

The schema for this algorithm is given on Figure S2.1(B) and, as in the main text, we assume a XY system where males are the heterogametic sex. The description follows the same structure as for sex-linked loci in the main text but note the three important methodological changes.

- The frequency of each mating type is calculated using the diploid genotypes of the brood survivors. This is given in the first line for each mating type (in the original formulation it was calculated from the gametic allele frequencies).
- Mutation occurs in the germline of the diploid brood survivors (Equation S2.1). This is simpler than the alternative approach of letting mutations occur in the mating types because each mating type would then have a larger number of brood genotype)
- Allele frequencies are extracted from diploid genotypes (Equation S1.2) rather than gametes

There are five diploid genotypes i.e.
♂(+Y), ♂(*m*Y), ♀(++), ♀(+*m*) ♀(*mm*)
Whose frequencies are denoted
f♂(+Y), f♂(*m*Y), f♀(++), f♀(+*m*), f♀(*mm*).

We store the frequencies of each diploid genotype after fRC in each of the six mating types in a matrix "freq_postRC" whose first element (row number) is the mating type number (see below) and the second element (column number) tracks each of the genotypes using the index:
♂(+Y) tracked as index 1
,♂(*m*Y) tracked as index 2
♀(++) tracked as index 3
♀(+*m*) tracked as index 4
♀(*mm*) tracked as index 5

*Mating type 1: ♂(+Y) with ♀(++)*
Frequency of this mating types is f♂*(+Y) * f♀(++)*
[Brood genotypes: 50% ♂(+Y), 50%♀(++)]

Size of brood after selection is $Z_1 = 1$
Size of brood after fRC is $B_1=1$

Frequencies of genotypes in broods after selection is (f)
f♂(+Y)= freq_postRC[1,1]= 0.5
f♂(*m*Y)=freq_postRC[1,2]=0
f♀(++)= freq_postRC[1,3]= 0.5
f♀(+*m*)= freq_postRC[1,4]= 0
f♀(*mm*)= freq_postRC[1,5]= 0

*Mating type 2: ♂(+Y) with ♀(+m)*
Frequency of this mating types is f♂*(+Y) *f♀(+m)*
[Brood genotypes: 25% ♂(+Y), 25% ♂(*m*Y), 25%♀(++), 25%♀(+*m*)]

Size of brood after selection is $Z_2$ = 0.25+ 0.25*(1-*s*)+ 0.25+ 0.25*(1-*h*s*)
Size of brood after fRC is $B_2=Z_2*C$ or $B_2=1$ whichever is the lower

Frequencies of genotypes in broods after selection is (f)
f♂(+Y)= freq_postRC[2,1]= 0.25/*N*
f♂(*m*Y)= freq_postRC[2,2]= 0.25*(1-*s*)/*N*
f♀(++)= freq_postRC[2,3]= 0.25/N
f♀(+*m*)= freq_postRC[2,4]= 0.25*(1-*h*s*)/*N*
f♀(*mm*)= freq_postRC[2,5]= 0
where *N* is a normalising factor equal to the sum of the numerators.

*Mating type 3: ♂(+Y) with ♀(mm)*
Frequency of this mating types is f♂*(+Y) *f♀(mm)*
[Brood genotypes: 50% ♂(*m*Y), 50%♀(+*m*)]

Size of brood after selection is $Z_3$ = 0.5*(1-*s*)+ 0.5*(1-*h*s*)
Size of brood after fRC is $B_3=Z_3*C$ or $B_3=1$ whichever is the lower

Frequencies of genotypes in broods after selection is (f)
f♂(+Y)= freq_postRC[3,1]= 0
f♂(*m*Y)= freq_postRC[3,2]= 0.5*(1-*s*)/*N*
f♀(++)= freq_postRC[3,3]= 0
f♀(+*m*)= freq_postRC[3,4]= 0.5*(1-*h*s*)/*N*
f♀(*mm*)= freq_postRC[3,5]= 0
where *N* is a normalising factor equal to the sum of the numerators.

*Mating type 4: ♂(mY) with ♀(++)*
Frequency of this mating types is f♂*(mY) * f♀(++)*
[Brood genotypes: 50% ♂(+Y), 50%♀(+*m*)]

Size of brood after selection is $Z_4$ = 0.5+ 0.5*(1-*h*s*)
Size of brood after fRC is $B_4=Z_4*C$ or $B_4=1$ whichever is the lower

Frequencies of genotypes in broods after selection is (f)
f♂(+Y)= freq_postRC[4,1]= 0.5/*N*
f♂(*m*Y)= freq_postRC[4,2]= 0
f♀(++)= freq_postRC[4,3]= 0
f♀(+*m*)= freq_postRC[4,4]= 0.5*(1-*h*s*)/*N*
f♀(*mm*)= freq_postRC[4,5]= 0
where *N* is a normalising factor equal to the sum of the numerators.

*Mating type 5: ♂(mY) with ♀(+m)*
Frequency of this mating types is f♂*(mY) *f♀(+m)*
[ Brood genotypes: 25% ♂(+Y), 25% ♂(*m*Y), 25%♀(+*m*), 25%♀(*mm*)]

Size of brood after selection is $Z_5$ = 0.25+ 0.25*(1-*s*)+ 0.25*(1-*h*s*)+ 0.25*(1-*s*)
Size of brood after fRC is $B_5=Z_5*C$ or $B_5=1$ whichever is the lower.

Frequencies of genotypes in broods after selection is (f)
f♂(+Y)= freq_postRC[5,1]= 0.25/*N*
f♂(*m*Y)= freq_postRC[5,2]= 0.25*(1-*s*)/*N*
f♀(++)= freq_postRC[5,3]= 0
f♀(+*m*)= freq_postRC[5,4]= 0.25*(1-*h***s*)/*N*
f♀(*mm*)= freq_postRC[5,5]= 0.25*(1-s)/*N*
where *N* is a normalising factor equal to the sum of the numerators.

*Mating type 6: ♂(mY) with ♀(mm)*
Frequency of this mating types is f♂*(mY) *f♀(mm)*
[Brood genotypes: 50% ♂(*m*Y), 50%♀(*mm*)]

Size of brood after selection is $Z_6$ = (1-*s*)
Size of brood after fRC is $B_6$=$Z_6$*C or $B_6$=1 whichever is the lower

Frequencies of diploid genotypes in broods after selection is (f)
f♂(+Y)= freq_postRC[6,1]= 0
f♂(*m*Y)= freq_postRC[6,2]= 0.5*(1-*s*)/*N*
f♀(++)= freq_postRC[6,3]= 0
f♀(+*m*)= freq_postRC[6,4]= 0
f♀(*mm*)= freq_postRC[6,5]= 0.5*(1-*s*)/*N*
where *N* is a normalising factor equal to the sum of the numerators.

We now calculate the number of each genotype in the next parent generation by summing the contribution from the six mating types, then normalise to convert to frequencies

$$\text{f}\male(+\text{Y}) = \frac{\sum_{i=1}^{6}(M_i * B_i * \text{freq\_postRC[i, 1]})}{N_m}$$

$$\text{f}\male(m\text{Y}) = \frac{\sum_{i=1}^{6}(M_i * B_i * \text{freq\_postRC[i, 2]})}{N_m}$$

$$\text{f}\female(++) = \frac{\sum_{i=1}^{6}(M_i * B_i * \text{freq\_postRC[i, 3]})}{N_f}$$

$$\text{f}\female(+m) = \frac{\sum_{i=1}^{6}(M_i * B_i * \text{freq\_postRC[i, 4]})}{N_f}$$

$$\text{f}\female(mm) = \frac{\sum_{i=1}^{6}(M_i * B_i * \text{freq\_postRC[i, 5]})}{N_f}$$

Where *i* is the mating type, $N_m$ is a normalising factor equal to the sum of the two numerators in the male genotype equations and $N_f$ is the corresponding female normalising factor equal to the sum of the three numerators in the female genotypes.

Mutation now occurs ignoring back mutation from mutant to wildtype and regarding the probability of a double mutation in the same individual as negligible. The genotype frequencies after mutation are denoted as $f'$.

Equations S1.1:

$$f' ♂(+Y) = f ♂(+Y) * (1-\mu)$$
$$f' ♂(mY) = f ♂(mY) + f ♂(+Y) * \mu$$
$$f' ♀(++) = f ♀(++) * (1-2\mu)$$
$$f' ♀(+m) = f ♀(+m) * (1-\mu) + f ♀(++) * 2\mu$$
$$f' ♀(mm) = f ♀(mm) + f ♀(+m) * \mu$$

Allele frequencies can then be extracted from these genotypes (either before or after mutation) using the standard symbols of *p* and *q* to represent wildtype and mutant allele respectively, and the subscripts *m* and *f* to denote males and females as

Equations S1.2

$$p_m = f' ♂(+Y)$$
$$q_m = f' ♂(mY)$$
$$p_f = f' ♀(++) + 0.5 * f' ♀(+m)$$
$$q_f = 0.5 * f' ♀(+m) + f' ♀(mm)$$

Note that that 2/3 of sex-linked alleles are in females, so overall frequencies of the mutant allele in the population, $\hat{q}$**, is**

$$\hat{q} = \frac{q_m}{3} + \frac{2 * q_f}{3}$$

Equations S1.3

Equilibrium allele frequencies was defined as occurring when the frequency of each of the five diploid genotypes different by a proportion less than 1±0.000001 in consecutive generations. As in the main text, the simulations were started with extremely low frequencies, and extremely high frequencies, and checked to confirm both iterate onto the same equilibrium value of $\hat{q}$ .

**References.**

1. Charlesworth B, Charlesworth D. Elements of evolutionary genetics. Colorado, USA: Roberts and company; 2010.

Figure S1.1. Schematic of the simulation. Panel (A) is the simulation tracking gametic allele frequencies, as in the main text. Panel (B) is the alternative simulation tracking allele frequencies in the diploid genotypes as described in this Supplementary Material. Note that "recombination" is used in its classic genetic sense of mixing of chromosomes which results in Hardy-Weinberg equilibrium within broods (Recombination #1) and populations (Recombination #2).

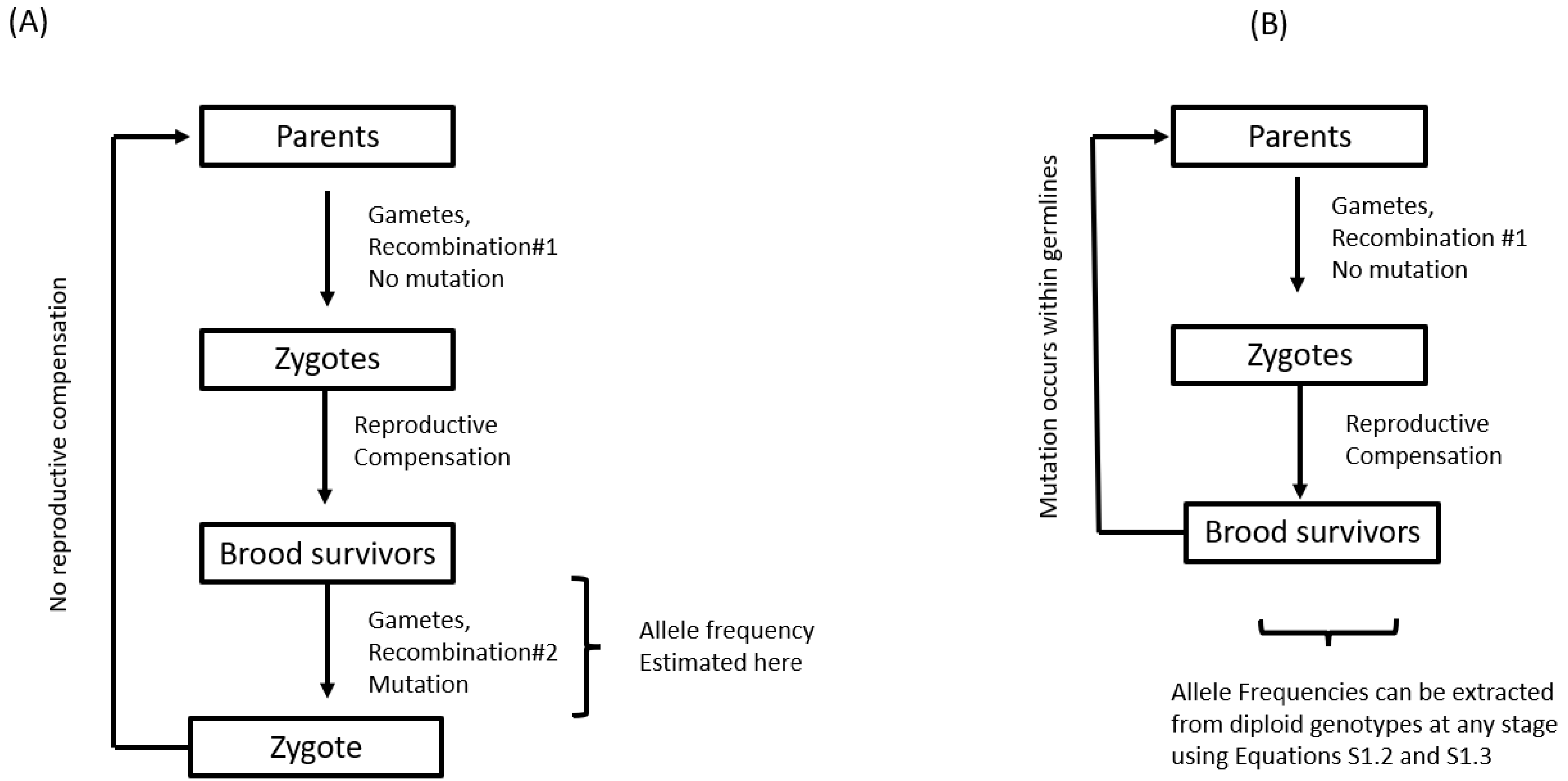

Figure S1.2. Anomalous results obtained for sex-linked loci when using the gametic allele frequencies algorithm (main text and Figure S1.1A) when tracking sexes separately and running males and female frequencies to equilibrium. The plot shows the increase in estimated equilibrium mutant allele frequencies assuming fRC is absent (i.e. with CR=1) compared with the textbook equation (i.e. Equation 7 of the main text). The results should be identical but diverge as selection coefficient increases and/or dominance decreases and may be up to 16% higher than the standard result when estimated using the algorithm.

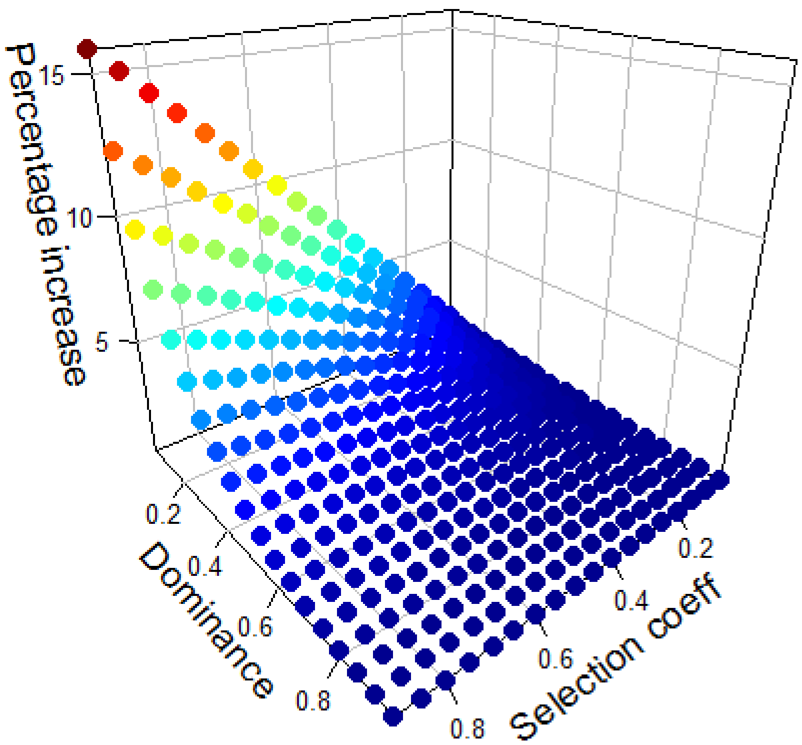
