## Supplementary Material, part 2 for "The impact of Fisher’s Reproductive Compensation on raising equilibrium frequencies of semi-dominant, non-lethal mutations under mutation/selection balance"

**Supplementary Material #2. Simulating the impact of fRC in species with complete sexual dimorphism.**

**1. Introduction.**

This Supplementary Material describes the impact of fRC on sex-linked genes in species that are sexually dimorphic to the extent that fRC can only occurs within sexes i.e. death of a male can only be compensated by increased survival of his brothers and not by increased survival of his sisters. Similarly for females i.e. death of female can only result in increased survival of her sister.

This was for intellectual curiosity, theoretical completeness, and to ensure the impact of fRC had been fully explored and was explicable across several demographic scenarios.

**2. Methods.**

The methodology is conceptually identical to the sex-linked calculations described in the main text i.e. assuming fRC acts of the whole brood. The only significant difference when considering sexual dimorphism is that fRC has to be calculated independently for each sex (so the maximum brood size of each sex is 0.5). The following description parallels that for sex-linked loci in the main text.

*Mating type 1: ♂(+Y) with ♀(++)*
Frequency of this mating types is $M_1$=f♀(+)* f♂(+)  ♀(+)
[Brood genotypes : 50% ♂(+Y), 50%♀(++)]

Size of male brood after selection and fRC  is  $B_1^m = 0.5$
Size of female brood after fRC and fRC is  $B_1^f = 0.5$

Frequencies of genotypes after selection within each sex is, respectively
f♂(+Y)=1
f♀(++)=1

And proportion of each gamete type transmitted from this mating type is
$P_1$♂(+)= f♂(=Y)
$P_1$♂(*m*)=0
$P_1$♀(+)= f♀(++)
$P_1$♀(*m*)=0

*Mating type 2: ♂(+Y) with ♀(+m)*
Frequency of this mating types is $M_2$= f♀(+)* [f♂(+)* ♀(*m*) + f♂(*m*)* f♀(+)]
[Brood genotypes : 25% ♂(+Y), 25% ♂(*m*Y), 25%♀(++), 25%♀(+*m*)]

Size of male brood after selection is $Z_2^m = 0.25 + 0.25 * (1 - s)$
Size of male brood after fRC is  $B_2^m = Z_2^m * \mathrm{C}$  or $B_2^m = 0.5$ whichever is the lower

Size of female brood after selection is $Z_2^f = 0.25 + 0.25 * (1 - h * s)$

Size of female brood after fRC is $B_2^f = Z_2^f * \mathrm{C}$ or $B_2^f = 0.5$ whichever is the lower

Frequencies of genotypes in broods after selection within each sex is (f)
f♂(+Y)=0.25/$N_m$
f♂($m$Y)=0.25*(1-$s$)/$N_m$
f♀(++)=0.25/$N_f$
f♀(+$m$)=0.25*(1-$h$*$s$)/$N_f$
where $N_m$ and $N_f$ are normalising factors equal to the sum of the numerators for the male and female gametes respectively.

Proportion of each gamete type transmitted from this mating type is
$P_2$♂(+)= f♂(+Y)
$P_2$♂($m$)= f♂($m$Y)
$P_2$♀(+)= f♀(++) + f♀(+$m$)*0.5
$P_2$♀($m$)= f♀(+$m$)*0.5

*Mating type 3: ♂(+Y) with ♀(mm)*
Frequency of this mating types is $M_3$=f♀(+)* f♂(m)* f♀(m)
Brood genotypes: 50% ♂($m$Y), 50%♀(+$m$)]

Size of male brood after selection is $Z_3^m = 0.5 * (1 - \mathrm{s})$
Size of male brood after fRC is $B_3^m = Z_3^m * \mathrm{C}$ or $B_3^m = 0.5$ whichever is the lower
Size of female brood after selection is $Z_3^f = 0.5 * (1 - \mathrm{h} * \mathrm{s})$
Size of female brood after fRC is $B_3^f = Z_3^f * \mathrm{C}$ or $B_3^f = 0.5$ whichever is the lower

Frequencies of genotypes in broods after selection within each sex is (f)
f♂($m$Y)=1
f♀(+$m$)=1

Proportion of each gamete type transmitted from this mating type is
$P_3$♂(+)= 0
$P_3$♂($m$)= f♂($m$Y)
$P_3$♀(+)= f♀(+$m$)*0.5
$P_3$♀($m$)= f♀(+$m$)*0.5

*Mating type 4: ♂(mY) with ♀(++)*
Frequency of this mating types is $M_4$= f♀($m$)* f♂(+)* ♀(+)
[Brood genotypes: 50% ♂(+Y), 50%♀(+$m$)]

Size of male brood after selection is $Z_4^m = 0.5$
Size of male brood after fRC is $B_4^m = Z_4^m * \mathrm{C}$ or $B_4^m = 0.5$ whichever is the lower
Size of female brood after selection is $Z_4^f = 0.5 * (1 - \mathrm{h} * \mathrm{s})$
Size of female brood after fRC is $B_4^f = Z_4^f * \mathrm{C}$ or $B_4^f = 0.5$ whichever is the lower

Frequencies of genotypes in broods after selection within each sex is (f)
f♂(+Y)=1

f♀(+$m$)=1

Proportion of each gamete type transmitted from this mating type is
$P_4$♂(+)= f♂(+Y)
$P_4$♂(m)= 0
$P_4$♀(+)= f♀(+$m$)*0.5
$P_4$♀(mut)= f♀(+$m$)*0.5

*Mating type 5: ♂(mY) with ♀(+m)*
Frequency of this mating types is $M_5$= f♀($m$)* [f♂(+)* f♀($m$) + f♂($m$)* ♀(+)]
[Brood genotypes: 25% ♂(+Y), 25% ♂($m$Y), 25%♀(+$m$), 25%♀($mm$)]

Size of male brood after selection is $Z_5^m = 0.25 + 0.25(1 - s)$
Size of male brood after fRC is $B_5^m = Z_5^m * \mathrm{C}$ or $B_5^m = 0.5$ whichever is the lower
Size of female brood after selection is $Z_5^f = 0.25 * (1 - \mathrm{h} * \mathrm{s}) + 0.25 * (1 - \mathrm{s})$
Size of female brood after fRC is $B_5^f = Z_5^f * \mathrm{C}$ or $B_5^f = 0.5$ whichever is the lower

Frequencies of genotypes in broods after selection within each sex is (f)
f♂(+Y)=0.25/$N_m$
f♂($m$Y)=0.25*(1-s)/$N_m$
f♀(+$m$)=0.25*(1-h*s)/$N_f$
f♀($mm$)=0.25*(1-s)/$N_f$
where $N_m$ and $N_f$ are normalising factors equal to the sum of the numerators for the male and female gametes respectively.

Proportion of each gamete type transmitted from this mating type is
$P_5$♂(+)= f♂(+Y)
$P_5$♂($m$)= f♂($m$Y)
$P_5$♀(+)= f♀(+$m$)*0.5
$P_5$♀($m$)= f♀(+$m$)*0.5+ f♀($mm$)

*Mating type 6: ♂(mY) with ♀(mm)*
Frequency of this mating types is $M_6$= f♀($m$)* f♂($m$)* f♀($m$)
[Brood genotypes: 50% ♂($m$Y), 50%♀($mm$)]

Size of male brood after selection is $Z_6^m = 0.5(1 - \mathrm{s})$
Size of male brood after fRC is $B_6^m = Z_6^m * \mathrm{C}$ or $B_6^m = 0.5$ whichever is the lower
Size of female brood after selection is $Z_6^f = 0.5 * (1 - \mathrm{s})$
Size of female brood after fRC is $B_6^f = Z_6^f * \mathrm{C}$ or $B_6^f = 0.5$ whichever is the lower

Frequencies of diploid genotypes in broods after selection within each sex is (f)
f♂($m$Y)=1
f♀($mm$)=1
.

Proportion of each gamete type transmitted from this mating type is
$P_6$♂(+)= 0
$P_6$♂(*m*)= f♂(*m*Y)
$P_6$♀(+)= 0
$P_6$♀(*m*)= f♀(*mm*)

Gamete frequencies are calculated as in the main text, noting the B (size of brood after fRC) is now separate for sexes i.e. "B" has superscripts 'm' or 'f' compared to equations in Section 2.3 of the main text

$$f'♂(+) = \frac{\sum_{i=1}^{6}\left(M_i * B_i^m * P_i♂(+)\right)}{N_m}$$

$$f'♂(\mathrm{m}) = \frac{\sum_{i=1}^{6}\left(M_i * B_i^m * P_i♂(\mathrm{m})\right)}{N_m}$$

$$f'♀(+) = \frac{\sum_{i=1}^{6}\left(M_i * B_i^f * P_i♀(+)\right)}{N_f}$$

$$f'♀(\mathrm{m}) = \frac{\sum_{i=1}^{6}\left(M_i * B_i^f * P_i♀(\mathrm{m})\right)}{N_f}$$

Where $N_m$ is a normalising factor equal to the sum of the two numerators in the male gamete equations and *Nf* is a normalising factor equal to the sum of the two numerators in the female gamete equations

Mutations were allowed to occur and equilibrium frequencies of mutant alleles were obtained as for sex-linked loci described in the main text.

**3. Results.**

When fRC is absent (obtained by setting CR=1) the algorithm for sexual dimorphism gives identical results to those obtained for sex-linked loci without polymorphism and recovers previous published results (i.e. Equations 7 and 8 of main text). The case of CR=1 generates the allele frequencies that occur in the absence of fRC so serve as the baseline scenario of non-fRC against which the impact of fRC is assessed.

Figure S2.1 is analogous to the results for autosomal and sex-linked loci given on Figure 2 of the main text; note the difference in the Y axis scale which, under sexual dimorphism is from 0 to 200% rather than from 0 to 100% on Figure 2 of the main text i.e. sexual dimorphism can potentially lead to fRC almost tripling equilibrium mutant allele frequencies. The relationship with *h*s* is given on Figure S2.2.

To understand the dynamics shown on Figure S2.1, recall that when the mutant allele is very rare most selection occurs in the following two mating types

Type 2: ♂(+Y) with ♀(+*m*)
Brood genotypes :25% ♂(+Y), 25% ♂(*m*Y), 25%♀(++), 25%♀(+*m*)

Type 4: ♂(*m*Y) with ♀(++)
Brood genotypes : 50% ♂(+Y), 50%♀(+*m*)

The pattern shown on Figure S2.1 is complicated and is best explained by considering the two cases of high and low dominance.

*(i) When dominance is low* (i.e. the mutation is largely recessive), most selection falls on the ♂ (*m*Y) male offspring which are only present in mating #2. Their death is largely compensated by their ♂(+Y) brothers i.e. removal of a mutant allele is usually compensated by increased survival of wildtype alleles. This makes fRC much less effective at raising equilibrium frequency of mutations compared to the absence of dimorphism (because, in the latter, death of ♂(*m*Y) in mating type #2 are partially compensated by increased survival of their ♀ (+*m*) sisters). Within this region of low dominance, selection acts as follows:

- At high values of *s*, most surviving brothers in the brood are ♂(+Y) so the loss of mutant alleles is almost entirely compensated by wildtype alleles, meaning fRC has little effect when the mutant has high selection coefficient and low dominance (bottom left hand corner of Figure S2.1 panels A and B).

- As the selection coefficient decreases, the proportion of ♂(*m*Y) among the surviving brothers increases so the impact of fRC increases i.e. death of mutant alleles tends to be compensated by survival of other mutant alleles in ♂(*m*Y) brothers; the impact of fRC consequently increases, and equilibrium frequency increases over the predictions of standard theory as shown along the left hand edge of Figure 2.1(A) and 2.1(B).

*(ii) As dominance increases*, more selection falls on the female ♀(+*m*) genotypes which occur in mating types 2 and 3. These female genotypes in mating type 2 are mostly compensated by + alleles from replacement ++ sisters which reducing the impact of fRC in this mating type. However, the impact of fRC in mating #4 will be much larger because only ♀(+*m*) are present among the female offspring, so their deaths are always compensated by survival of the same genotype and selection against the mutant allele in females in this brood is

effectively absent until fRC can no longer fully compensate for these female deaths. When CR=1.5 full compensation can occur until >33% of the female brood die (Figure 1 of main text) so the lines representing s= 0.5, 0.6, 0.8 and 0.9 in FigureS2.1(B) shows a declining impact of fRC once $h*s$ is greater than 33% i.e. when $h$ exceeds 0.67, 0.56, 0.42, 0.37 respectively for these values of s.

Figure S2.2 shows the relationship between mutant equilibrium frequency and the magnitude of $h*s$. The interplay between dominance and selection coefficient described above means the relationship is far more complicated than for autosomal or non-dimorphic sex-linkage (Figures 3 and 4 respectively of the main text) and I was unable to find an algebraic solution for the equilibrium frequencies. As for sex-linked loci in non-dimorphic species (main text) a linear regression of the increases obtained for CR=20 was implemented by fitting $s$, $h$ and $s*h$ and returned the empirical result that the fold-increase compared to the standard result in the absence of fRC, ω, is

$\omega = 2.16 + 0.94h - 0.96s - 0.45hs$ Equation S2.1

or, with negligible impact on accuracy

$\omega = 2.16 + 0.95(h - s) - 0.45hs$ Equation S2.2

These empirical estimates lie within a proportion 0.96 to 1.13 of the exact results with the higher deviations at low values of dominance and, to a lesser extent high values of selection coefficient. Reducing parameter space to vary dominance from 0.2 to 0.8 and selection coefficient from 0.1 to 0.7 results in

$\omega = 2.2 + 0.90h - 0.95s - 0.44hs$ Equation S2.3

or, with negligible impact on accuracy

$\omega = 2.2 + 0.925(h - s) - 0.44hs$ Equation S2.4

With values of ω within the range 0.98 to 1.05 of the simulation values. Recall. however, that fRC is far less effective at restoring brood size under assumption of sexual dimorphism, so the approximation breaks down at high levels of $h*s$ when fRC is unable to replace the dead brood members of the same sex (e.g. Figure S2.1B).

In summary, fRC within sexual dimorphism can potentially increase equilibrium frequency above standard theory to a much greater extent than autosomal loci or sex-linkage without sexual dimorphism i.e. generating a maximum increase of ω≈3 compared of a maximum increase of ω≈2 in the other two scenarios.

Figure S2.1. Percentage increase of equilibrium mutant allele frequency attributable to fRC acting on sex-linked loci in the presence of complete sexual dimorphism (i.e., compared to frequency in absence of fRC). Panel (A) shows the relationship with selection coefficient and dominance, assuming CR=1.5 and mutation rate is $10^{-5}$. Panel (B) shows transects across the plot shown in Panel (A)

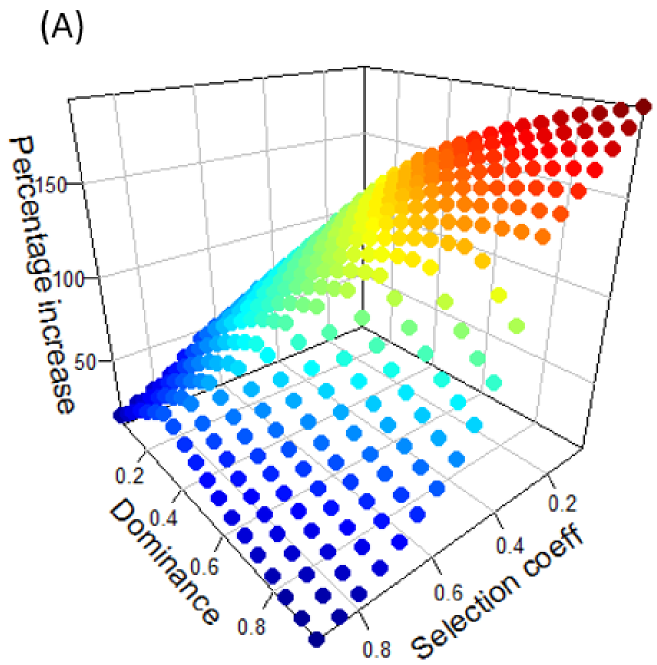


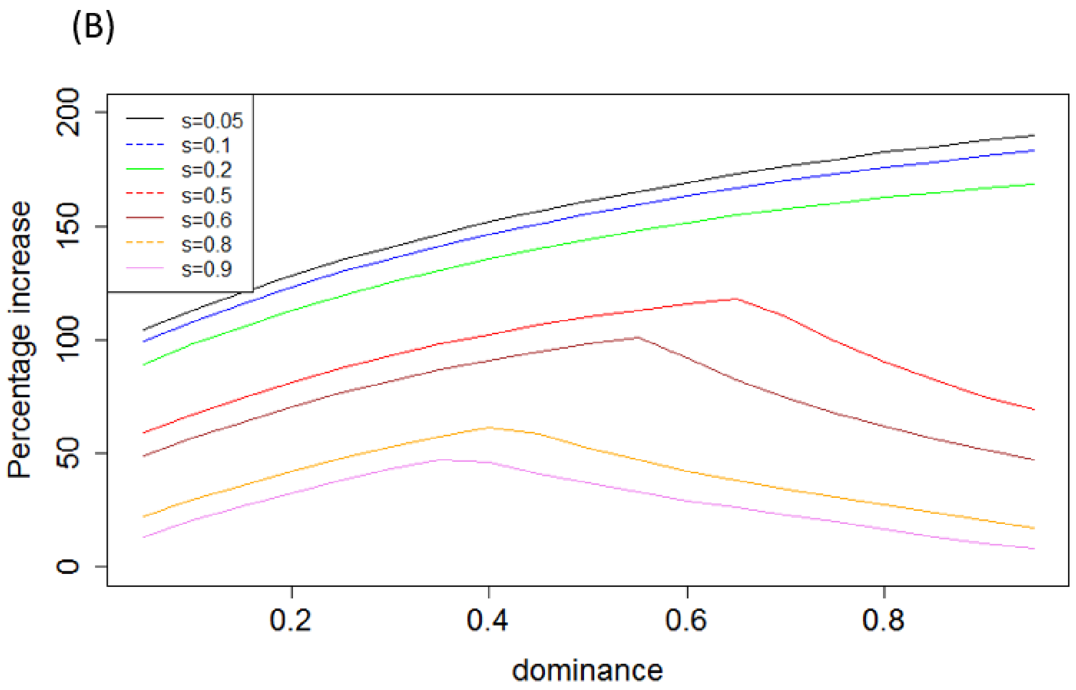

Figure S2.2. Panels (A) and (B) are as for Figure 3 of the main text but for sex-linked loci with fRC acting within sexual dimorphism with values for CR=1.2 and CR=1.5.

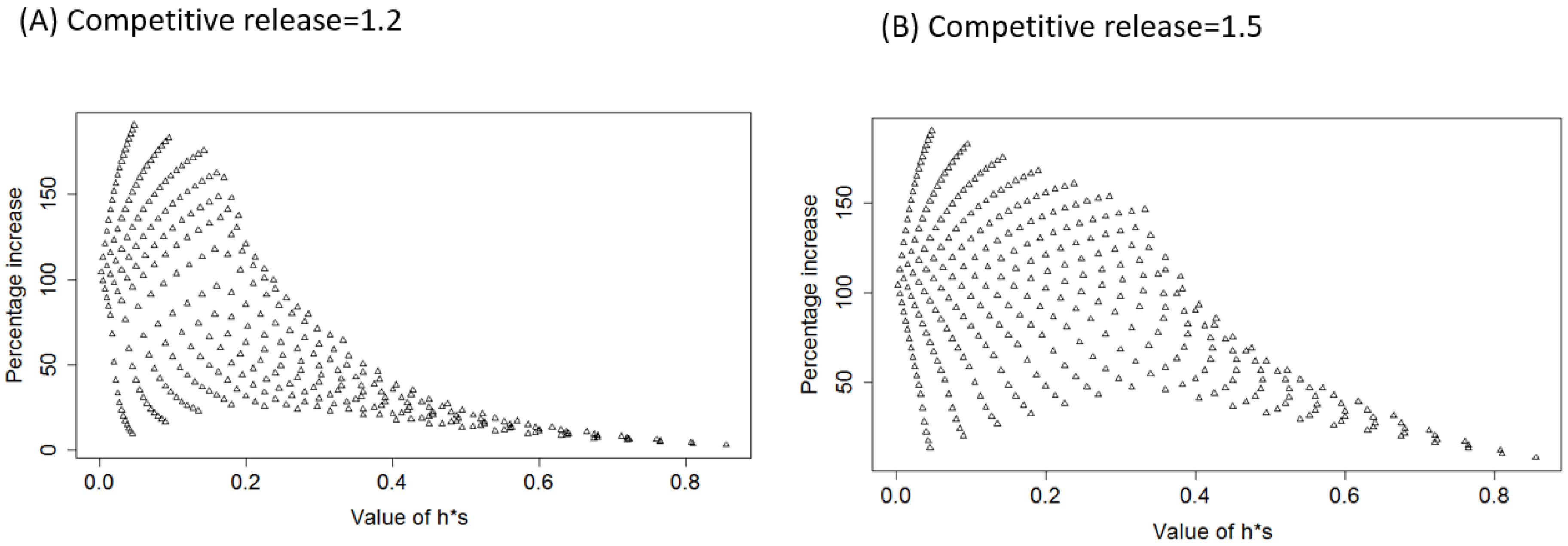
